# Systematic discovery of single-cell protein networks in cancer with Shusi

**DOI:** 10.1101/2025.04.27.649905

**Authors:** Tianyun Zhang, Jiajun Yu, Shang Lou, Yaozhen Liang, Yue Liang, Zhekai Li, Haishuai Wang, Shanshan Pei, Ning Shen

## Abstract

Context-specific protein-protein interaction (PPI) drive heterogeneity of primary tumor, forming a formidable challenge to effective cancer therapy. However, systematically mapping and modeling these interactions at single-cell resolution across diverse cancer types remains an unmet need. Here, we present Shusi, a large language model-enhanced variational graph auto-encoder model trained on over single cells across 23 cancer types, to predict context-specific PPIs. Shusi outperforms existing state-of-the-art methods, as validated through orthogonal experimental evidence. Cancer-specific mutations are significantly enriched in Shusi-predicted networks, offering complementary insights to conventional marker gene-based approaches. Through systematic evaluations, we demonstrate strong associations between Shusi-predicted network topologies, genetic vulnerabilities, and therapeutic sensitivity. Finally, in acute myeloid leukemia (AML), a blood cancer where cell-state heterogeneity drives clinical resistance, Shusi pinpointed potential targets as actionable vulnerabilities of resistant leukemia subpopulations, as validated experimentally in primary AML. Shusi offers a deep-learning tool for implementing precision medicine based on context-specific protein network architecture.

## Introduction

Cancer cells exhibit highly context-specific interactions and signaling cascades, reflecting extensive inter- and intra-tumor heterogeneity^1–5^. This heterogeneity drives cancer initiation, progression^6,7^, and the development of drug resistance^8,9^. Consequently, drug development strategies are evolving from targeting isolated molecular factors to targeting critical cell populations, opening new avenues for precision medicine^10^. The core objective of precision oncology is to effectively target and eliminate malignant cells across diverse tumor subpopulations^11,12^. Consequently, strategies to dissect this heterogeneity have been a long-standing interest but remain challenging^11,13^.

Single-cell RNA sequencing (scRNA-seq) enables large-scale transcriptional profiling of individual cells, offering high-resolution insights into cellular systems^9,14^. Within these systems, gene transcription responds to intracellular and extracellular signals to coordinate cellular activities. Gene co-expression networks derived from transcriptomic data offer a significant framework for modeling gene–gene interactions and functional relationships^15–17^, with broad applications in characterizing disease states^18^ and identifying potential drug targets^19^. However, coexpression networks cannot distinguish between direct and indirect regulatory relationships and cannot reflect downstream functions.

Integrating scRNA-seq with multi-omics data provides opportunities to expand the universe of cancer targets^20,21^. Proteins, as fundamental functional units^22^, offer critical insights into phenotype-genotype relationships. Although AI-driven advances enable protein structure prediction^23^, function annotation^24^, and target identification^25,26^, these tools predominantly depend on static PPI databases that fail to capture in vivo context-dependent interactions. Recent efforts to develop context-aware PPI predictions through multimodal deep learning^26–28^ represent significant progress. However, AI frameworks capable of integrating scRNA-seq with dynamic PPI networks to enable precision cancer therapeutics remain unrealized. To address this gap, we introduce a deep learning framework that synergizes single-cell transcriptomics with protein interaction landscapes to identify clinically actionable drug targets.

We introduce Shusi, named after a therapeutic eagle-like divine bird from the ancient Chinese legend “Classic of Mountains and Sea”. Shusi is a context-aware variational graph auto-encoder (VGAE)^29^ model that integrates large language models (LLMs)^30^ for network analysis. By leveraging single-cell transcriptomics alongside protein interaction networks, Shusi generates contextualized interactions, producing 5,198,624 contextualized protein interactions across 22,722 proteins within 71,575 pan-cancer networks. Notably, Shusi accurately predicted cell-type-specific networks that aligned with experimental findings in relevant cancer contexts. By exploring various networks across 23 cancer types, we demonstrate the model’s ability to identify context-specific essential genes and nominate potential therapeutic targets through network analyses. Finally, applying Shusi to fine-tune on scRNA-seq data from acute myeloid leukemia (AML) patients, we identified novel actionable therapeutic vulnerabilities of therapy-resistant leukemia, validated through experiments on primary AML samples. This approach directly addresses the clinical challenge of developmental heterogeneity in AML. Being adaptable to various operational context, Shusi has the potential to pave the way for the widespread application of foundational models tailored to diverse biological landscapes.

## Results

### The Shusi model

The design principle of Shusi is to leverage scRNA-seq data with PPI information. This approach is designed to elucidate gene-to-gene relationships acorss cells (Fig 1a).

**Fig. 1.**
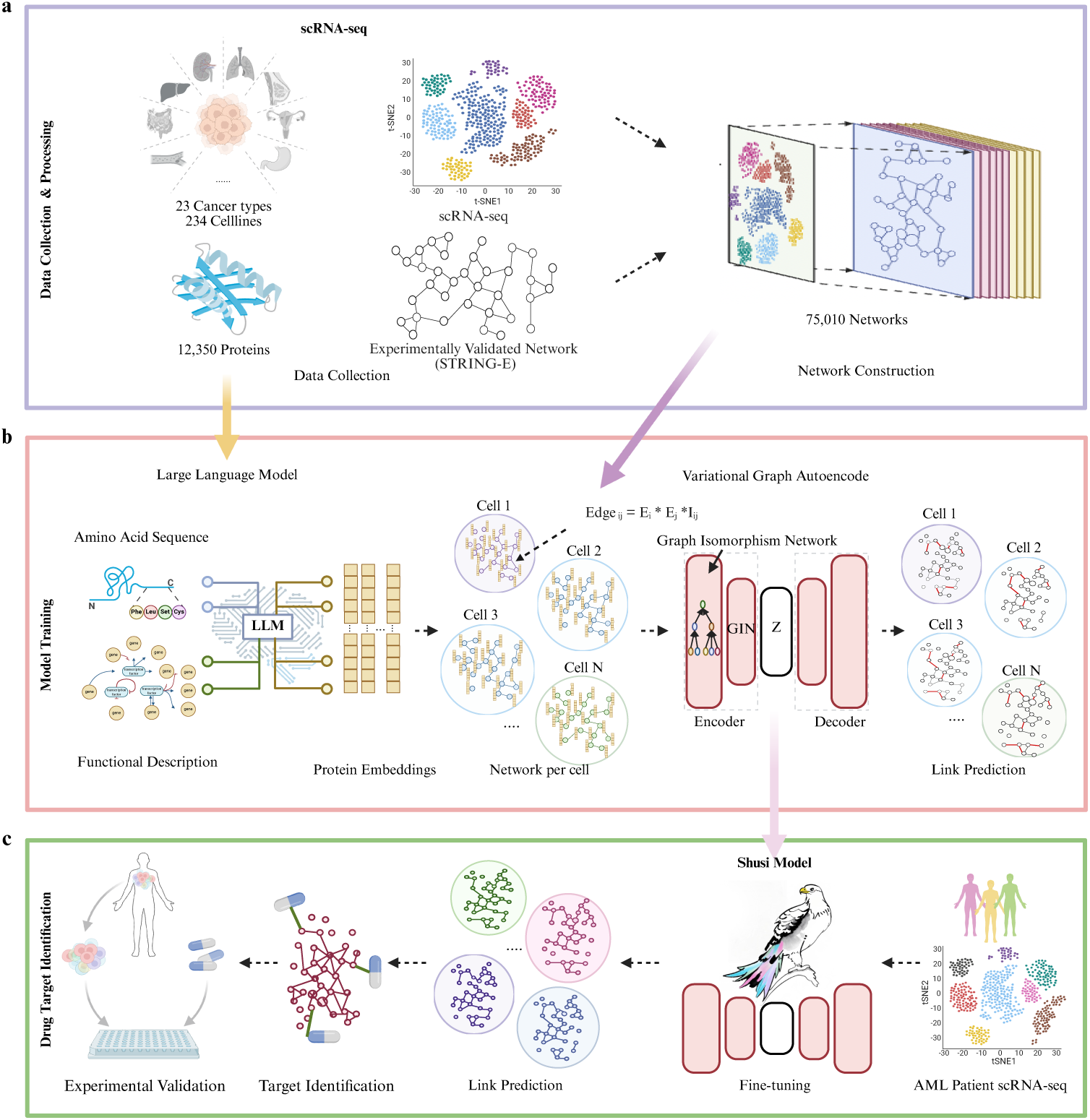
Overview of Shusi. a, Pipeline for constructing single-cell protein interaction networks from scRNA-seq data. b, Architecture of the Shusi model for predicting novel interactions on single cell level. d, Fine-tuning and application of Shusi to AML patient scRNA-seq data

To enable cell-specific interaction prediction, we integrated 75,010 single-cell transcriptomic profiles from 23 cancer types, generated using two complementary sequencing platforms^31,32^, the experimental-validated STRING database (STRING-E) as reference PPI network^33^(Fig. 1a, Fig. 2a). The reference interactome comprised 22,722 protein-coding genes curated from the STRING database (v11.0)^33,34^, providing experimentally validated interactions, functional annotations, and amino acid sequences. We derived single-cell-specific networks containing an average of 3,681 proteins connected by about 496,179 high-confidence interactions for model training (Methods). This approach allowed us to capture biological heterogeneity at superior resolution, enabling precise identification of context-dependent interactions. Meanwhile, our single-cell network construction approach ensured that the model could handle variable input sizes, accommodating any number of cells while maintaining computational scalability.

**Fig. 2.**
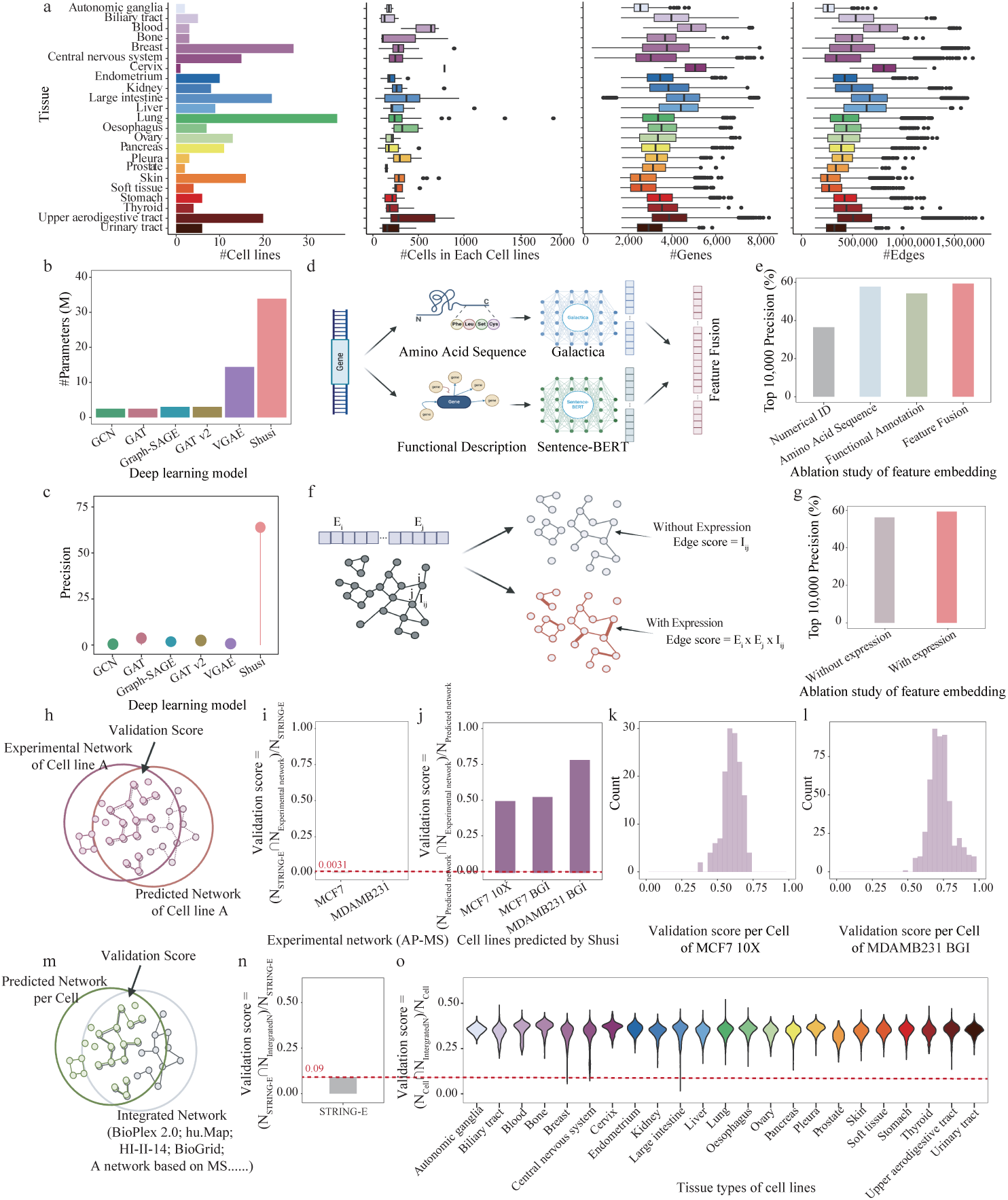
Evaluation of Shusi’s performance. a, Datasets used for model training. The bar plot shows the distribution of cell lines across 23 cancer types. Box plots (left to right) depict: (i) the number of cells per cell line, (ii) the number of genes per training network, and (iii) the number of edges per training network. b, Comparison of model parameters between Shusi and baseline methods. c, Precision of edges predicted by Shusi and baseline models compared to experimentally validated interactions in STRING-E. d, Workflow of the ablation study evaluating node embedding contributions. e, Top 10,000 precision scores from the node embedding ablation study. f, Workflow of the ablation study evaluating edge embedding contributions. g, Top 10,000 precision scores from the edge embedding ablation study. h. Validation scores between novel edges predicted by Shusi and experimentally detected interactions. i, Validation scores comparing interactions in STRING-E with experimentally detected interactions. j, Validation scores comparing Shusi-predicted interactions with experimentally detected interactions. k, l, Cell-specific validation scores comparing Shusi-predicted interactions with experimentally detected interactions for MCF7 (k) and MDAMB231 (l) cell lines. m, Validation scores between novel edges predicted by Shusi and interactions in the integrated network. n, Validation scores comparing interactions in STRING-E with those in the integrated network. o, Cell-specific validation scores comparing Shusi-predicted interactions with interactions in the integrated network.

Shushi learned both the functions and sequences of proteins by integrating two large language models (LLMs) to encode proteins to construct the node embeddings in the graph (Fig. 1b, Fig. S1a, Methods). The dual-encoding strategy processed amino acid sequences and functional descriptions separately. Sentence-BERT^35^ effectively encoded amino acid sequences, similar to how text is embedded in LLMs. Whereas the functional descriptions were encoded using Galactica^36^, an LLM pre-trained on 48 million scientific documents. This dual approach enables the model to simultaneously capture sequence-derived properties and functional roles, enhancing its ability to predict biologically relevant interactions.

For network modeling, Shusi combines a variational graph auto-encoder (VGAE)^37^ with a graph isomorphism network (GIN) in a symmetric encoder-decoder structure(Fig. 1b, Fig. S1a-b). Unlike conventional approaches restricted to static network analysis, our model dynamically processes over 71,000 single-cell-specific biological networks, each representing distinct cellular conditions, by leveraging the scalability of VGAE for large graph datasets. The encoder employs a dual-layer GIN structure that analyzes adjacency matrices combined with dual-encoded node features (sequence and functional embeddings), generating compact latent representations that retain context-sensitive topological relationships. Thus, the enhanced discriminative power, derived from the Weisfeiler-Lehman graph isomorphism framework, enables precise detection of nuanced topological variations across cellular contexts, such as cell-specific interaction patterns. For interaction reconstruction, the decoder calculates pairwise similarity scores between latent node representations using inner products and applies sigmoid activation to predict interactions (Methods).

Post-training, Shusi generated novel, context-specific protein interactions for each cell, capturing the heterogeneity of protein networks. In the following sections, we validate the meaning and accuracy of single-cell context-specific novel networks predicted by Shusi. We show that the single-cell networks capture cell-specific patterns and accurately reconstruct biologically relevant connections. Moreover, we demonstrate that the reconstructed single-cell networks offer orthogonal insights into exploring essential genes within cells (Fig. 1c).

### Performance evaluations of Shusi

We systematically evaluated Shusi’s performance through benchmark analysis against baseline models. To ensure comparability, we randomly partitioned 15,010 single cells (20% of the dataset) as an independent test set, masking 20% of protein interactions in each cellular network for prediction. Baseline models—averaging 2.5M parameters (excluding VGAE at 14.39M)—exhibited comparable performance (Fig. 2b). In contrast, Shusi (33.88M parameters) demonstrated significantly superior predictive accuracy, achieving the highest values across all performance metrics(Fig. S2).

To establish an optimal edge-ranking threshold for practical applications, we systematically evaluated thresholds corresponding to the top 1,000, 10,000, 20,000, and 50,000 predicted edges (Fig. S3). Analysis revealed that the 10,000-edge threshold provides maximal stability across datasets with variable cell counts—a critical factor for scRNA-seq data robustness. We therefore recommend this threshold as the standard for downstream applications.

We further validated the contributions of both dual-encoder feature extraction module and the construction of context-specific networks to Shusi’s superior performance through an ablation study (Fig. 2d, e). To quantify feature contributions, we systematically ablated components by retraining Shusi with alternative embeddings: numerical IDs (baseline), amino acid sequences (Sentence-BERT), and functional descriptions (Galactica). The complete model achieved peak performance with a precision of 0.59 (Fig. 2e), confirming the necessity of integrating structural and functional information. Additionally, weighted graphs further enhanced interaction prediction accuracy compared to unweighted graphs (Fig. 2f, g). Thus, the ablation study validates the advantages of our node feature embedding and network construction strategy.

Next, we evaluated Shusi’s ability to predict novel interactions across independent experiments. We compared these predictions to cancer-specific PPIs identified through an independent high-throughput affinity-purification mass spectrometry (AP-MS) experiment focused on breast cancer cell lines MCF7 and MDAMB231^38,39^ (Fig. 2h). Notably, Shusi-predicted interactions showed a validation score of 60.67% with AP-MS-derived PPIs, which is significantly higher than the 0.23% and 0.31% consistency rates observed between STRING-E and AP-MS for MCF7 and MDAMB231, respectively (Fig. 2i-l). This suggests that Shusi captured higher proportions of cell-specific interactions compared with the prior context-free database. At the single-cell level, validation scores exhibited minor variations, further supporting Shusi’s ability to capture cell-specific information (Fig. 2k, l; Fig. S2a).

To extend our analysis to a broader context, we benchmarked Shusi’s outputs against several static protein interaction databases. We found 8.8% of Shusi-predicted interactions received multi-evidence support (text mining, co-expression, co-occurrence) beyond STRING’s experimentally validated interactions (Fig. S2b). We next compiled an integrated database of cancer-specific experimental PPIs and data from five existing protein association databases (Methods, Fig. 2m)^25^. Compared to the integrated database, Shusi demonstrated a 3.8-fold higher validation score against STRING-E (34.40% vs. 9.07%; Fig. 2n, o), validating the reliability of Shusi predicted interactions. Additionally, single-cell resolution analysis revealed stable performance across all cancer types and cell lines (Fig. 2o), further validating Shusi’s robustness. These results demonstrate Shusi’s capacity to resolve cellular diversity and identify context-dependent interactions that may not be captured in static databases.

### Shusi-informed proteins are enriched for cancer-specific mutations

The analysis of scRNA-seq typically focuses on identifying marker genes through differential expression, which limits subsequent biological findings to explore interactions across genes.

Thus, we hypothesized that edges and nodes predicted by Shusi may provide orthogonal biological insights beyond those derived from marker gene-based analysis.

Shusi encompassed 71,575 cell-specific networks covering 22,722 proteins and generated 5,198,623 novel interactions (Fig.3a, b). These networks exhibited variations in both edges and nodes. To systematically explore this diversity, we calculated node degrees by summarizing connected edges for each gene to represent network connections (Fig. S3a). By clustering node degrees for each cell and visualizing their distributions using t-distributed stochastic neighbor embedding (tSNE) colored by cancer types (Fig 3c), we found that cells tended to group closer with those of the same cancer type (Fig. S3b). While node degrees offered insights into the importance of nodes within network structures, we also investigated interaction similarities. Across 234 cell lines, we discovered approximately 349,209 cell-type-specific interactions. The diversity of interactions appeared somewhat influenced by the cell population of each cell line (Fig. S3c). Additionally, we noted higher interaction consistency rates within the same cancer type (Fig. S3c), which confirms that Shusi predicted networks align with the intra-tumor and inter-tumor heterogeneity biology. Therefore, the single-cell networks predicted by Shusi underscore the heterogeneity exhibited by different cancer types based on various node degrees and edge connections in their network structures.

**Fig. 3.**
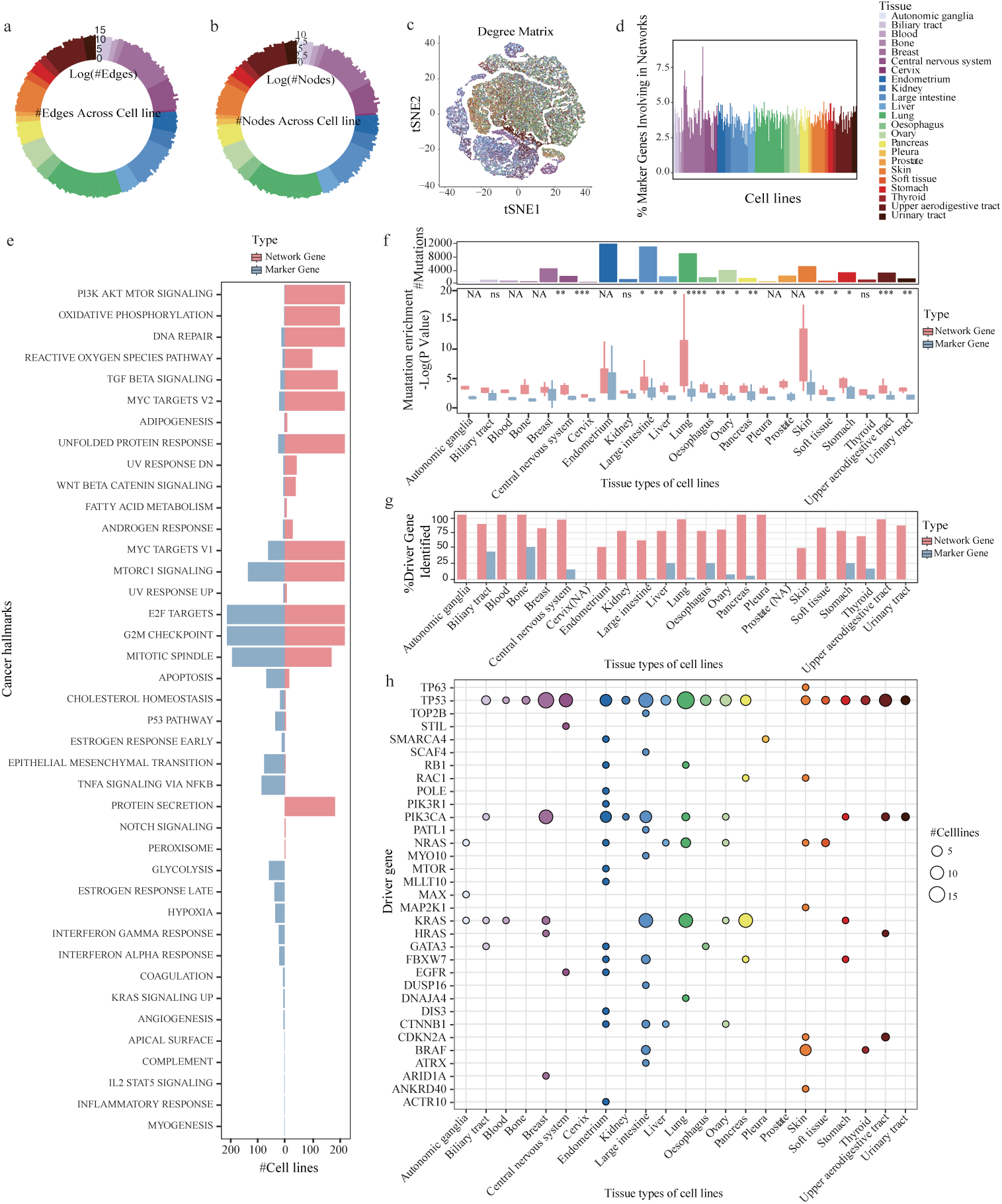
Evaluation of novel edges detected by Shusi. a, Bar plot shows the number of edges in Shusi-predicted novel networks across cell lines. b, Bar plot shows the number of nodes in Shusi-predicted novel networks across cell lines. c, UMAP visualizes the degree matrix across single cells, highlighting cell-type-specific network topologies. d, Bar plot shows the percentage of marker genes included in Shusi-predicted novel networks. e, Enrichment analysis of cancer hallmarks for nodes in Shusi-predicted novel networks compared to marker genes. f, Top: Bar plot shows the number of mutation-harboring genes per cancer type. Bottom: Boxplot compares the enrichment of mutation-harboring genes in Shusi-predicted novel networks versus marker genes across cancer types. NA represents that the sample size is less and equal to three. g, Comparison of the percentage of driver genes detected in Shusi-predicted novel networks versus those identified as marker genes. NA represents that no driver gene has been recorded in the CCLE database. h, List of driver genes identified in Shusi-predicted novel networks across 23 cancer types.

By incorporating novel networks derived from cell lines, our analysis uncovered approximately 3,530 proteins engaged in potential interactions within specific cell-type contexts (Fig 3b). We then assessed network-related genes to ensure they delineate distinct network structures rather than solely reflecting fluctuating gene expression levels similar to marker genes. Among the total gene pool of 22,722, 3.75% were designated marker genes in the same cell line (Fig 3d). The remaining genes, previously unexplored in disease contexts, demonstrated independence from marker genes of the same cell line, often exhibiting lower expression levels (Fig. S4a,b). Network genes demonstrated significantly broader functional relevance than marker genes, with enrichment in cancer Hallmark pathways observed across substantially more cell lines. Importantly, these network genes revealed specific mechanistic associations with oncogenic pathways, including Notch signaling, peroxisome regulation, and protein secretion, providing new molecular insights into cancer pathogenesis (Fig 3e).

The characterization of cancer genomes has yielded fundamental insights into somatically altered genes across tumor types, transforming our understanding of cancer biology and enabling their identification as therapeutic targets^10^. As our framework relies exclusively on scRNA-seq without genomic sequencing, we investigated whether this approach could provide insight into these somatically altered genes to inform tailored therapeutic strategies. Consequently, we compared our framework’s performance against conventional differential gene expression (DGE) analysis for identifying these somatically altered genes.

We analyzed the somatic mutations for each cell line sourced from the DepMap database^40^ to probe the enrichment of such mutations within the novel interactions predicted by Shusi. Our analysis identified 953,941 PPIs involving 5,022 proteins that host at least one disease-associated somatic mutation. Notably, the majority of these interactions (88.66%) exhibited mutations on one protein, while the remainder (11.34%) showcased mutations on both interacting proteins. This finding emphasize Shusi-predicted PPIs on the impact of disease-associated mutations.

To further evaluate the enrichment of disease-associated mutations within genes prioritized by Shusi, we conducted a comprehensive comparison against differentially expressed genes (DEGs) across 234 cell lines. Strikingly, Shusi identified an average of 84 mutant genes per cell line, compared to only 10 somatically altered genes detected by DEGs. Critically, Shusi-predicted genes showed significant enrichment for somatic mutations versus marker genes across all 23 cancer types—independent of overall mutation burden (Fig. 3f, S5). Furthermore, driver mutations—functionally crucial for oncogenesis—were captured at substantially higher rates by Shusi’s network analysis than by conventional marker gene approaches (Fig. 3g). These driver mutations exhibited cancer-type specificity, with canonical alterations like BRAF V600E prominently enriched in melanoma (Fig. 3h). Collectively, Shusi’s novel interaction predictions provide biologically orthogonal insights into cancer mechanisms, powerfully complementing traditional DEG analysis.

### Novel networks accurately reflect cellular perturbation response

Identifying essential genes for cancer cell survival represents a promising therapeutic strategy. To evaluate whether our novel predicted networks reflect functional gene essentiality without relying on external screening data, we leveraged genomewide CRISPR-Cas9 viability data from DepMap—encompassing 11,763 protein-coding genes across 186 cancer cell lines (Methods, Fig. 4a)^40^. This independent dataset provides a rigorous benchmark to validate the biological significance of Shusi-predicted networks and demonstrate their capacity for recommending therapeutic targets.

**Fig. 4.**
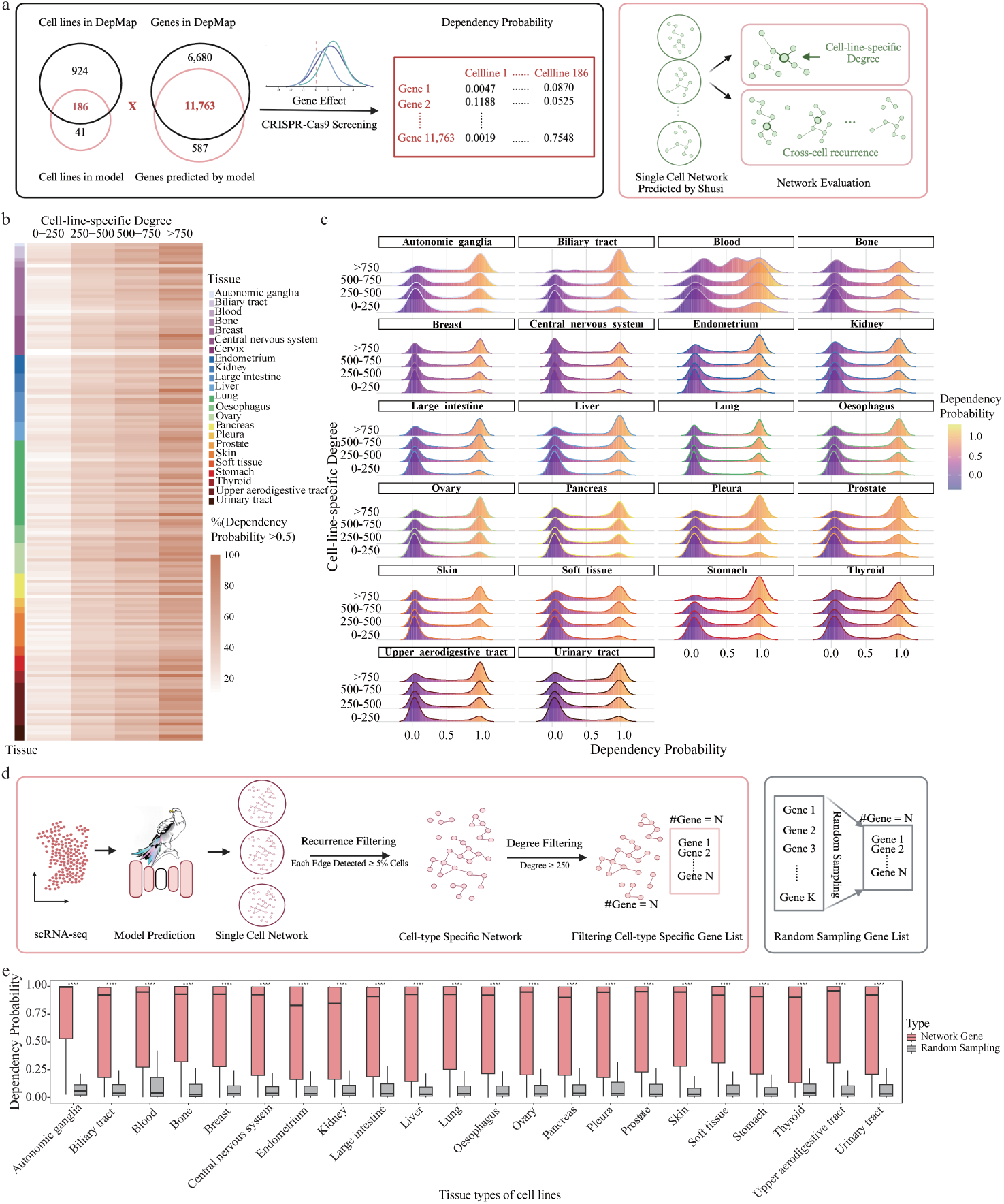
Evaluation of dependency probabilities for nodes in novel networks. a, Workflow illustrates the collection of dependency probabilities from the CCLE and the computation of node degrees and cross-cell recurrence ratios based on network topologies. b, Heatmap shows the percentage of genes with dependency probabilities > 0.5 across different degree centrality groups. c, Distribution of dependency probabilities across different degree centrality groups. d, Pipeline for selecting the filtered cell-type-specific gene list and the randomly sampled gene list. e, Boxplot comparing dependency probabilities between the filtered cell-type-specific gene list and the randomly sampled gene list.

We quantified gene essentiality using two network topological metrics: cross-cell recurrence (frequency of gene detection across single-cell networks) and degree centrality (summed interaction weights in cell-type-specific networks) (Methods, Fig. 4a). Cross-cell recurrence measures pan-contextual regulatory influence, defined as the proportion of cells wherein a gene participates in functional modules. Degree centrality reflects network integration strength.

We classified genes into five groups based on their recurrences and assessed the percentage of genes with a dependency probability greater than 0.5 within each group. This stratification by recurrence revealed a striking 3.10-fold enrichment of essential genes (dependency probability *>* 0.5) from lowest (20%) to highest recurrence groups (61.9%) (Fig. S6a). Low-recurrence genes (¡20% frequency) exhibited baseline dependency (*µ* = 0.22), while high-recurrence genes demonstrated therapeutic vulnerability (*µ* = 0.60) (Fig. S6b), establishing recurrence as a robust feature of biological necessity.

By consolidating cell-line-specific networks across all 186 cancer models, we computed the degree for each gene (Methods, Fig 4a). Degree-stratified analysis revealed a strong positive correlation between connectivity and essentiality. Genes in the highest connectivity quintile (*>* 750 edges) demonstrated 83.6% essentiality (dependency probability *>* 0.5), compared to 12.3% in the low-connectivity group (*<* 250), suggesting the significance of network connectivity to evaluate cellular fitness.

Implementing a dual-threshold filter (edge recurrence ≥ 5% and degree centrality ≥ 250), we identified 11,324 high-confidence targets (median 93 genes/cell line). These targets exhibited significantly elevated dependency probabilities compared to background genes (*µ* = 0.67 vs *µ* = 0.15) (Fig 4e). The robustness of this approach was further evidenced by conserved topology-essentiality relationships; genes with higher levels of cross-cell recurrences and degrees demonstrated greater therapeutic vulnerability, enabling the identification of actionable targets. These findings establish network topology as a powerful instructor of genetic vulnerabilities, providing a framework to prioritize therapeutic targets.

### Pharmacogenomic landscape of Shusi-predicted interactions

To systematically investigate the relationship between novel networks and therapeutic vulnerability, we integrated context-aware interaction networks with drug sensitivity profiles from the Genomics of Drug Sensitivity in Cancer (GDSC) database^41^. Our analysis encompassed 1,000 cancer cell lines exposed to 263 therapeutic compounds, focusing on 143 cell lines with 15 clinically actionable compounds targeting 51 ChEMBL-annotated genes (Fig. 5a).

**Fig. 5.**
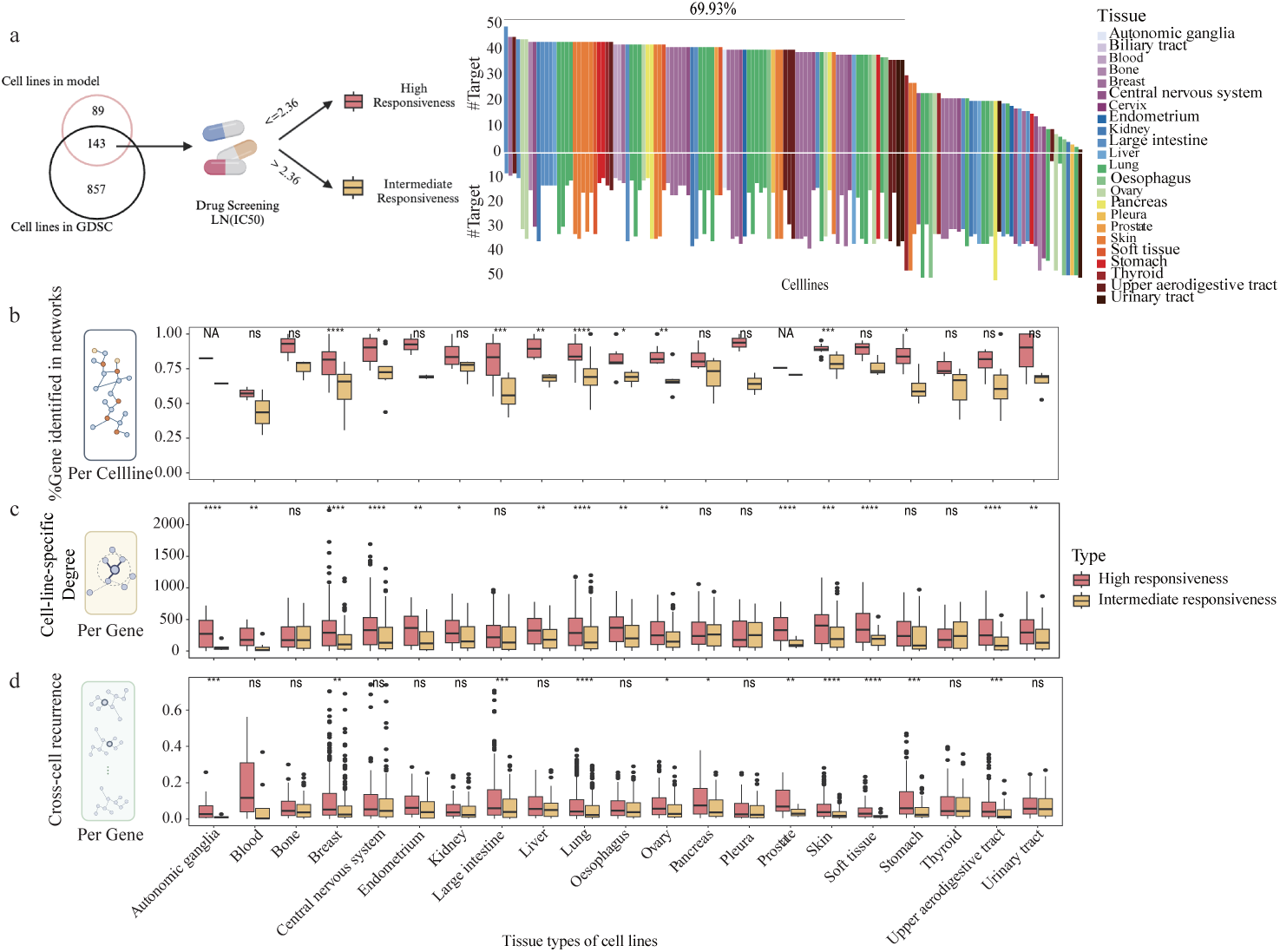
Performance of identifying drug target genes in novel networks. a, Workflow for classifying drugs into high- and intermediate-responsiveness groups for each cell line. The bar plot shows the number of drug target genes for high- and intermediate-responsiveness drugs across cell lines. b, Boxplot comparing the percentage of high- and intermediate-responsiveness drug targets identified in Shusi-predicted networks. c, Boxplot comparing the degree centrality of high- and intermediate-responsiveness drug targets in Shusi-predicted networks. d, Boxplot comparing the cross-cell recurrence of high- and intermediate-responsiveness drug targets in Shusi-predicted networks.

Using natural log-transformed IC50 values, we stratified compounds into high- and intermediate-responsiveness groups of each cell line (Methods). High-responsiveness compounds targeted significantly more genes per cell line, with 69.9% of cell lines containing over 30 high-responsiveness target genes (Fig. 5a).

Next, we constructed cell-line-specific networks by consolidating newly identified interactions across all cells (Methods), and explored the presence of high-responsiveness drug targets within these cell-line-specific networks. Our analysis demonstrated a greater coverage of high-responsiveness drug targets compared to intermediate-responsiveness targets within our network (Fig 5b). This finding suggests that when searching for potential drug target genes through novel networks, there is a higher likelihood of identifying high-responsiveness drug targets.

We further investigated the relationship between network characteristics and drug responses. Our analysis revealed that target genes with higher degrees or greater cross-cell recurrences exhibited significant drug sensitivity within cell lines, in contrast to genes associated with intermediate drug responses (Fig. 5c,d). This relationship was particularly pronounced in autonomic ganglia and prostate cancer cell lines, where intermediate-responsiveness targets showed markedly reduced network integration (Fig 5c, d). This divergence suggests that topological matrices can be used to explore target vulnerability across cellular contexts.

### Shusi uncovers actionable vulnerabilities of therapy-resistant leukemia disease

AML is a clinically aggressive hematologic malignancy presenting extensive inter-and intra-patient cell-state heterogeneity across the myeloid developmental hierarchy^4,5,42–44^, offering an ideal model for implementing Shusi to develop therapeutic regimens. To enable precise prediction on primary AML samples, we first fine-tuned Shusi using scRNA-seq profiles from 32,146 cells across 27 primary AML patients^45^. Through systematic analysis of protein interaction networks predicted by Shusi, we delineated the topological architectures of AML at single-cell resolution (Fig. 6a). Our tSNE analysis, focusing on the degree matrices derived from single-cell networks, revealed that primary AML cells, residing in distinct cell states across the myeloid developmental hierarchy, exhibited different distribution patterns of degree matrices (Fig. 6b, Fig. S7a-g). Quantitative analysis of cell-state-specific interactomes revealed an average of 5,832 genes and 1,158,259 interactions per state (Fig. S8a, b), with only 2,869 genes and 292,443 edges conserved across all seven myeloid cell states, reflecting the significant protein network heterogeneity present among them.

Importantly, previous studies, including our own, have shown that primitive (Prim) subsets, including Leukemia Stem and Progenitor Cell (LSPC)-quiescent, LSPC-primed, and LSPC-cycling, preferentially rely on the antiapoptotic protein BCL-2 for survival thus exhibit exclusive sensitivity to small molecule BCL-2 inhibitor venetoclax in the clinic. In contrast, monocytic (Mono) subsets encompassing Promonocyte (Pro-Mono)-like and Monocyte (Mono)-like cell states are resistant to BCL-2 inhibition and can contribute to relapse and refractory disease, forming a formidable challenge in the AML clinic^5,45,46^. As shown in Fig. 6c-d, distribution of degree matrices is vastly different between the Venetoclax-sensitive Prim and Venetoclax-resistant Mono subsets, provoking the need to identify novel therapeutic targets specific to the Mono subsets based on their unique protein network. Applying Shusi, we discovered that the Mono-specific protein network includes 759 nodes and 9,950 edges while excluding edges shared with Prim networks (Fig. 6e). Pathway enrichment analysis revealed that the Mono-specific network enriches for TNF-*α* signaling, TGF-*β* signaling, allograft rejection and others, consistent with studies reporting inflammation and immune evasion as characteristics of monocytic leukemia^4,47,48^ (Fig. 6e, Fig. S8c).

Leveraging these network topologies, we explored their utility in predicting Mono-specific drug targets. According to analysis in Fig. 4, degree of connectivity is a strong predictor of biological dependency. Therefore, we implemented a stringent topological selection criterion, applying a top 10% connectivity threshold as a cut-off (Fig. 6f), when analyzed Shusi-derived Mono-specific networks (Methods, Fig. S11a), This systematic approach identified 75 Mono-specific targets, providing a reservoir of plausible targets to eradicate this venetoclax-resistant disease population (Supplementary Table 1).

Finally, to validate the predictions, we built an in vitro drug sensitivity assay coupled with multi-color flow cytometry-based immunophenotyping analysis, allowing evaluation of targeting efficacy at disease subpopulation level. We performed this analysis on a representative primary AML specimen with mixed monocytic and primitive (MMP) disease subpopulations (Fig. 6g, h, Fig. S9a), as described in our previous work^45^. Notably, we verfied that these Shusi-predicted targets were over-expressed in various genetic subtypes of human AML relative to normal hematopoietic stem and progenitor cells, suggesting their plausible therapeutic potential (Fig. S10). In conclusion, these results demonstrated the exciting potential of Shusi in designing novel effective drug combinations to address the clinically challenging developmental heterogeneity of human cancer.

## Discussion

We introduce Shusi, a transformative model that bridges single-cell transcriptomics with context-aware interactome modeling. By integrating 75,010 single-cell protein interaction networks across 23 cancer types, Shusi achieves unprecedented single-cell resolution in predicting cellular context-dependent interactions. The dual LLM architecture enables Shusi to integrate both functional annotations and amino acid sequence information for comprehensive gene characterization. Complementing this, the VGAE with GIN architecture empowers Shusi to extract global topological features from single-cell networks. Together, these computational frameworks allow the model to learn core protein interaction patterns to predict novel cell-specific interactions (Fig. S1b cell 1 vs cell 2). Large-scale statistical analyzes highlight the significance of the network structures predicted by Shusi, revealing a significant enrichment of somatic cancer mutations within these interactions. This approach not only deepens our understanding of protein interactions in disease but also provides a powerful tool for drug discovery and therapeutic development. Comprehensive analyzes demonstrate Shusi’s ability to identify potential drug target genes, offering profound insights for therapeutic innovation. To facilitate widespread adoption and further development, we have implemented Shusi as a user-friendly software package, enabling the broader scientific community to conduct systematic analyzes and build upon our model.

Traditional context-free protein-protein interaction networks are inherently limited in their ability to capture the dynamic and context-dependent nature of protein interactions across diverse cell types and tissues. Shusi addresses this limitation by generating context-specific interaction networks tailored to individual cells. Our results show that these contextual networks significantly enhance the understanding of protein roles by accounting for functional variability across different cellular environments. These cell-specific networks enable the identification of therapeutic candidates taking into account intra-tumor heterogeneity. Future progress in understanding protein functions and developing targeted therapies will require mapping both cell-type-specific protein roles and their context-dependent interactions that frequently occur beyond canonical protein complexes. Models like Shusi are poised to play a pivotal role in realizing this potential by predicting single-cell novel protein networks. These networks can then be used to identify therapeutic targets, advancing the field of precision medicine.

Human cancer is marked by heterogeneity of cell-states deviated from normal developmental trajectory^3^. Acute myeloid leukemia (AML), as an example, is a clinically aggressive hematologic malignancy presenting extensive inter- and intra-patient cell-state heterogeneity across the myeloid developmental hierarchy^4,5,42^. The success of the therapeutic regimen nominated by Shusi demonstrates it’s unique advantage in identifying actionable vulnerabilities of cell-state-specific cancer. Interestingly, the drug targets that successfully kills the monocytic leukemia disease overlap rarely with genes in the most enriched pathways we identified for Mono-specific network. They however all present ample connections to the enriched pathways (Fig. 6e), suggesting that key therapeutic targets accounting for vulnerabilities of cell-state-specific cancer may not directly lie in the most enriched pathways. However, through integration of the context-specific protein network predicted by Shusi and in vitro validation assays in primary AML samples, we validated the therapeutic potential of targeting genes in combination with the commonly used Ven/Aza regimen to treat developmentally heterogeneous AML disease. Relapse and refractory responses to Venetoclax are significant challenges in the field of AML^49^. The discovery of these vulnerabilities enabled by Shusi warrants further investigation in preclinical models.

Our study acknowledges two key limitations in contemporary interactomics. First, the incomplete human reference interactome limits ground-truth validation^50^. Second, while single-cell transcriptomic integration mitigates context-blindness, network inference may introduce false positives. Despite these challenges, Shusi’s performance—validated through CRISPR screening and GDSC-tested compound screening—demonstrates that integrating cellular context is not merely beneficial but essential for target discovery. Future iterations will incorporate spatial proteomics and single-cell proteomic data to resolve subcellular interaction dynamics. Shusi’s architecture is designed to integrate these data, creating a virtuous cycle between prediction and experimental validation.

Shusi exemplifies how context-aware modeling can decode drug targets. By mapping these dynamic interaction landscapes, we move closer to realizing the promise of precision medicine. Therapies are tailored to the ever-changing interaction networks that define cellular identity. As single-cell multi-omics datasets continue to expand, models like Shusi will be critical for transforming this data deluge into actionable biological insights.

## Methods

### Data collection and preprocessing

#### Reference human physical protein interaction network

The reference PPI network was constructed using experimentally validated interactions from the STRING database (STRING-E) (version 11.0)^33^. To ensure high-confidence interactions, we implemented stringent filtering criteria retaining only those physical associations supported by experimental evidence.

#### Single-cell RNA sequencing of cancer cell lines

We utilized two comprehensive single-cell transcriptomic datasets for model training: the pan-cancer atlas from Kinker et al. (GSE157220)^31^, and the multi-omics profiling data from Zhu et al. (https://db.cngb.org/cdcp/scatlashcl/download/)^32^. The Kinker dataset comprises single-cell RNA sequencing (scRNA-seq) profiles of 198 cancer cell lines spanning 22 distinct cancer types, and the Zhu dataset provides the transcriptomic characterization of 40 cancer cell lines. We obtained a combined dataset of 71,575 single-cell profiles corresponding to 232 unique cancer cell lines. We collected differentially expressed genes detected in these two studies for further analysis.

#### Single-cell RNA sequencing of AML clinical samples

ScRNA-seq data were obtained from 27 primary AML specimens, as previously described by Pei et al. (GSE232559)^45^. The dataset comprised 32,416 high-quality single-cell transcriptomes, representing 14 distinct cell populations across the myeloid differentiation hierarchy.

#### Construction of single cell networks

To establish context-aware single-cell protein interaction networks, we integrated gene expression profiles with prior knowledge of protein-protein interactions. For each pairwise gene combination, the edge weight *W_ij_* was computed as:

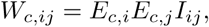

where *E_c,i_* and *E_c,j_* represent the normalized expression levels of gene i and gene j in cell c, respectively, and *I_i_j* denotes the experimentally validated interaction weight from the STRING database. This integrative approach ensures that edge weights reflect both transcriptional co-expression patterns and physical interaction probabilities, enabling the construction of cell-specific interaction landscapes.

Within each individual cell, these weights undergone normalization to mitigate potential biases introduced by variations in sequencing technologies and data processing pipelines:

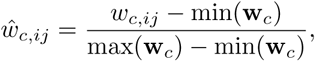

where *ŵ_c,ij_* ∈ R represents the normalized edge weight between genes *i* and *j* in cell *c*, and **w***_c_* ∈ R^|*E*|×1^ denotes the full set of edge weights in cell *c*, and |*E*| refers to the number of the edges in cell *c*.

### Model Framework

The **Shusi** model is composed of three primary modules: the feature extraction module, the encoder, and the decoder. The feature extraction module is responsible for extracting features of genes including amino acid sequences and functional annotations. The encoder maps these features into a latent space, while the decoder reconstructs the likelihood of interactions.

#### Feature Extraction Module

The feature extraction module operated on amino acid sequences and functional annotations collected from the STRING database. To simultaneously model both the sequence and text information, we utilized two distinct encoders: Sentence-BERT^35^ and Galactica^36^. Sentence-BERT (SBERT) is a transformer-based model trained on masked token prediction tasks, designed to generate semantic sentence representations. On the other hand, Galactica is a large language model specifically trained on scientific texts, optimized for next-token prediction tasks. Let the gene annotation be denoted by *a* and *s*, where *a_i_* represents the textual annotation for gene *i*, and *s_i_* represents the gene sequence.

The embeddings for each cell were obtained by concatenating the embeddings generated by both encoders. Specifically, for gene *i*, the combined embedding 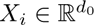 was given by:

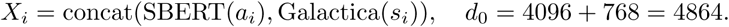

Here, *X_i_* denoted the concatenated embedding for gene *i*, with a total dimension of 4864.

#### Encoder

After obtaining the embeddings 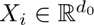 for all genes and the corresponding edge weights *ŵ_c,ij_* ∈ R for all edges in cell *c*, we applied a GIN^29^ within a VGAE^37^ framework. The initial embeddings at layer 0 were set as *h*^0^ = *X_i_*. The update rule for each subsequent layer of the GIN was defined as:

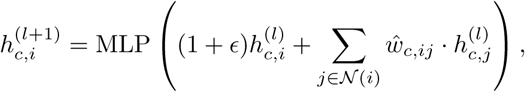

where:

- *ɛ* ∈ R is a learnable parameter.
- 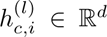 represents the embedding of gene *i* at layer *l* for cell *c*, with hidden dimension *d*.
- N (*i*) is the set of neighbors of gene *i* in cell *c*.
- *ŵ_c,ij_* ∈ R is the normalized weight of the edge between gene *i* and gene *j* at layer *l* in cell *c*.
- MLP refers to a multi-layer perceptron applied to aggregate the information from neighbors.

Furthermore, to capture additional edge information, we introduced an edge encoder, which projected edge weights to a feature space of the same dimension as the node embeddings, i.e., *e_c,ij_* = *W ŵ_c,ij_*), where *e_c,ij_* ∈ R*^d^* and *W* ∈ R*^d^*^×1^. The node embedding 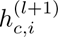 was then updated as:

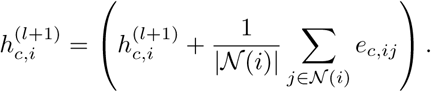

The message passing process was defined as *g_ϕ_* (*X_i_,* **w***_c_*) for gene *i* and the graph weight information with *ϕ* denotes to all the learnable parameters. For the VGAE framework, node embeddings were passed through two layyers to generate the mean *µ_c,i_* ∈ R*^d^* and the standard deviation log *σ_c,i_* ∈ R*^d^* for the latent variable of gene *i* in cell *c*:

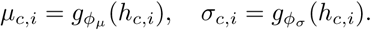

The latent variable *z_c,i_* ∈ R*^d^* for gene *i* in cell *c* was then represented as:

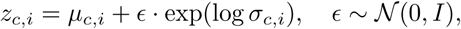

where *ɛ* ∈ R*^d^* is a random variable drawn from a standard normal distribution.

#### Decoder

The decoder estimated the likelihood of an edge between two genes in cell *c* by computing the inner product of their latent variable embeddings, followed by a sigmoid activation to obtain the final connection probability:

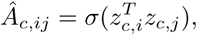

where:

- *Â_c,ij_* ∈ [0, 1] is the predicted probability of an edge between genes *i* and *j* in cell *c*.
- *σ*(·) is the sigmoid function, which maps the inner product to a probability value.

#### Loss Function

The loss function for the VGAE was a link prediction loss, which aims to predict the presence of edges between genes based on their latent variable embeddings. Binary cross-entropy loss was used to compute the loss between the predicted edge probabilities and the actual observed edges. Given the predicted probability *Â_c,ij_* and the ground truth adjacency matrix *A_c,ij_* for cell *c*, the link prediction loss was defined as:

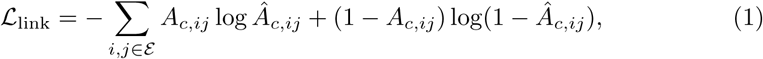

where: - *A_c,ij_* ∈ {0, 1} is the ground truth adjacency matrix, where *A_c,ij_* = 1 if there is an edge between genes *i* and *j*, and 0 otherwise. - *Â_c,ij_* = *σ*(*z^T^ z_c,j_*) is the predicted probability of an edge between genes *i* and *j*, computed using the inner product of their latent embeddings followed by a sigmoid activation.

The total loss function combined the link prediction loss with the regularization terms on the latent variable distributions. The final objective was:

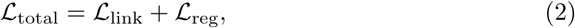

where *L*_reg_ is a regularization term that encourages the latent variable distributions to be close to a standard normal distribution.

### Network prediction benchmarking

#### Baseline models

GCN is a graph-based convolutional network that learns node representations by aggregating weighted information from neighboring nodes^51^. It leverages the graph Laplacian matrix and its normalized form for feature propagation and local smoothing. GraphSAGE generates node embeddings by sampling a fixed number of neighbors and aggregating their features^52^. Unlike GCN, it operates efficiently on large-scale graphs by focusing on local neighborhoods and supports inductive learning for unseen nodes.

GAT incorporates attention mechanisms to assign different importance weights to each node’s neighbors^53^. By adaptively learning the contribution of neighbors, GAT enhances the flexibility of feature aggregation and improves performance on heterogeneous graphs.

GAT v2 is an enhanced version of GAT, introducing dynamic attention in place of static attention^54^. This improvement makes weight computation more sensitive to input features, enhancing its expressive power and enabling GATv2 to better capture complex relationships between nodes.

VGAE extends the variational autoencoder framework to graph data for unsupervised representation learning^37^. By learning the latent distribution of nodes, VGAE excels in tasks such as link prediction and node classification, while also supporting generative modeling of graphs.

#### Network prediction task and experimental setting

Network prediction aims to infer missing edges between nodes in a graph based on observed graph structure and node features. For our benchmarking experiments, we established a rigorous evaluation framework to assess model performance. For baseline models, we utilized a direct training approach without pre-training. We partitioned the cell graph edges with 80% allocated for training and propagation, while reserving 20% for evaluation. Within the training portion, we further divided the edges, with 80% used for node information aggregation (effectively 64% of all edges) and the remaining 20% (16% of all edges) dedicated to parameter optimization. The held-out 20% of edges served as the test set for performance evaluation.

For Shusi, our comprehensive dataset consisted of 75,010 cell graphs. We randomly shuffled these graphs and designated 80% for pre-training, while using the remaining 20% for evaluation—maintaining consistency with the evaluation set used for baseline models. For each training cell graph, we employed a similar edge allocation strategy, using 80% of edges for information propagation and 20% for parameter updates. For the evaluation cell graphs, we likewise allocated 80% of edges for propagation and reserved 20% for performance assessment.

Regarding model configurations, baseline models utilized 32-dimensional ID embeddings as initial node features and were optimized with a learning rate of 1e-3. In contrast, Shusi employed a learning rate of 5e-4 during pre-training and leveraged rich, biologically-informed representations by concatenating features from Galactica and Sentence-BERT, resulting in 4864-dimensional initial node features. All models incorporated weight decay of 1e-5 and featured two layers of GNN propagation with a consistent hidden dimension of 512 to ensure fair comparison. Besides, both the baseline model and Shusi are built using the PyTorch^55^ and PyG^56^ frameworks, and both utilize the Adam optimizer^57^.

### Ablation study

To systematically evaluate the impact of different input features on Shusi’s performance, we conducted an ablation study across 60,000 cell graphs. First, we initialized node embeddings using random numerical IDs as a baseline. Next, we independently tested the contributions of amino acid sequence embeddings (generated by Galactica) and functional annotation embeddings (derived from Sentence-BERT). To assess the influence of edge features, we compared model performance between two configurations: (1) interactions incorporating gene expression profiles *W_ij_* = *E_i_E_j_I_ij_* and (2) interactions without expression weighting *I_ij_*.

### Fine-tuning Shusi for AML datasets

After pre-training Shusi on 80% of our comprehensive single cell graph dataset, we implemented a targeted fine-tuning approach for AML applications. We fine-tuned Shusi using scRNA-seq profiles from 32,146 cells across 27 primary AML specimens (GSE232559).

Our fine-tuning methodology maintained the architectural integrity of the pre-trained model while adapting it to AML-specific cellular contexts. For each AML cell graph, we allocated 80% of edges for information propagation and reserved the remaining 20% for parameter updates. To preserve the stability of Shusi’s core parameters while allowing sufficient adaptation to AML-specific features, we employed a reduced learning rate of 1e-5 during fine-tuning process.

The fine-tuning process retained representations established during pre-training, continuing to leverage the concatenated features from Galactica and Sentence-BERT that yield 4864-dimensional node features. We maintained the weight decay parameter at 1e-5 and preserved the two-layer GNN propagation architecture with a hidden dimension of 512.

### Metrics and statistical analyses

Here, we described metrics, visualization methods and statistical tests used in our analysis.

#### Validation scores compared with gold-standard reference networks

To systematically evaluate the reliability of novel interactions predicted by Shusi, we established a multi-tier validation framework. For MCF7 and MDAMB231 cell lines, we curated PPI datasets generated through affinity purification coupled with mass spectrometry (AP-MS)^39^, representing gold-standard reference networks. For cell lines lacking matched experimental PPI data, we employed the integrated protein interaction repository compiled by Zheng et al.^25^, which aggregates high-confidence physical interactions from multiple orthogonal studies including BioGRID^58^, BioPlex^59^, hu.Map^60^, Hein et al.^61^ and HURI^50^.

The network consistency was quantified using the validated rate metric:

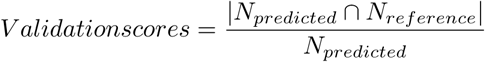

where *N_predicted_* represents edges in Shusi-generated networks and *N_reference_* denotes interactions in the reference networks.

#### Visualization of degrees

In the consolidated network, the single cell network degree (*D_g_*) of each gene was computed by summing all connected edges:

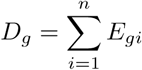

where *E_gi_* represents edges between gene *g* and its interacting partners.

To visualize the topological properties of Shushi-predicted networks, we performed dimensionality reduction using t-distributed stochastic neighbor embedding (tSNE). Network degree distributions were processed using Seurat (v5.1.0)^62^ in R, with single-cell network degrees transformed into two-dimensional embeddings while maintaining default parameter settings.

#### Euclidean distance

Based on the tSNE embeddings, we quantified both intra-cancer and inter-cancer Euclidean distances to assess cellular heterogeneity. For intra-cancer distances, we calculated pairwise Euclidean distances using the dist function with the Euclidean method in R. For inter-cancer distances, we evaluated all possible pairs among the 23 cancer types (totaling 253 unique combinations). The Euclidean distance between two cancer types *i* and *j* was computed as:

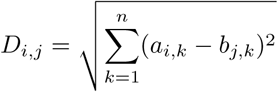

where (*a_i,k_* and *b_j,k_* represent the tSNE coordinates of cells from cancer types *i* and *j* respectively. For each cancer type, we derived the average Euclidean distance by computing its mean distance to the remaining 22 cancer types.

#### Enrichment analysis

Enrichment analysis was conducted on both marker genes and network-predicted genes using the clusterProfiler package (version 4.12.6) in R. Specifically, we performed Hallmark enrichment analysis from the MSigDB(v7.5.1) to identify biological pathways significantly associated with these genes.

#### Statistical tests

For two-group comparisons, we employed Student’s t-test. Analyses with fewer than three biological replicates per group were excluded from statistical evaluation and annotated as NA (not applicable) in subsequent results.

Cell line-specific somatic mutation profiles were obtained from the DepMap (24Q4). To evaluate the biological significance of mutated genes, we conducted comparative enrichment analyses between two gene sets: (1) network-predicted genes and (2) cell type-specific marker genes.

Using a hypergeometric test framework, we calculated the statistical overrepresentation of mutated genes against genome-wide background expectations with the following parameters:

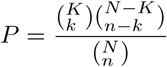

where *N* = total protenin-coding genes, *K* = mutated genes in cell line, *n* = gene set size (network/marker genes), *k* = overlapping mutated genes.

### CRISPR-Cas9 Screeing analysis

Gene dependency probabilities were obtained from the DepMap (24Q4)^40^. We established the pharmacogenomic dataset by systematically matching cell line identifiers between DepMap and our study cohort. The final analytical matrix encompassed quantitative dependency profiles for 11,763 protein-coding genes across 186 cancer cell lines.

For each cell line, we quantified the cross-cell recurrence of each gene, defined as its frequency of occurrence across all corresponding single-cell networks (Fig. S4). This metric was calculated as:

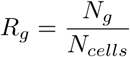

where *R_g_* = recurrence ratio of gene g, *N_g_*= number of single-cell networks containing gene g, and *N_cells_*= total single-cell networks per cell line.

We integrated single-cell networks at the cell line level by aggregating edges across individual cells. In the resulting cell-line-specific network, an edge is retained if it appears in at least one constituent single-cell network, with edge multiplicity consolidated to binary presence (present/absent) regardless of its occurrence frequency across cells. In the cell-line-specific network, the cell-line-specific degree (*D_g_*) of each gene was computed by summing all connected edges:

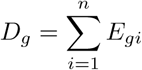

where *E_gi_* represents edges between gene *g* and its interacting partners (Fig. S4).

To systematically prioritize therapeutic targets, we developed a network-based filtering pipeline that integrates topological features to refine candidate gene lists. The pipeline implements two sequential filters: (1) Edge persistence filter: Retaining interactions detected in *>* 5% of single-cell networks within each cell line to ensure context-specific reproducibility; (2) Connectivity threshold: Selecting genes exceeding a cell-line-specific degree threshold of 250, based on prior topological analysis demonstrating this cutoff optimally distinguishes essential hubs. For rigorous benchmarking, we conducted matched random sampling of genes equivalent in number to network-filtered candidates within each cell line. This comparative framework enables quantitative evaluation of network-guided prioritization efficacy against null expectations.

### Drug sensitivity data

Drug response data comprising natural logarithm-transformed half-maximal inhibitory concentration (Ln(IC50)) values for all screened cell line-drug combinations were obtained from the GDSC^41^. Drug target annotations were retrieved from ChEMBL (https://www.ebi.ac.uk/chembl/)^63^ using standardized chemical identifiers (ChEMBL IDs). Through systematic matching of cell line identifiers (COSMIC IDs) between GDSC and our study cohort, we established a unified pharmacogenomic dataset. The final analytical matrix encompassed 143 cancer cell lines across 15 therapeutic compounds targeting 52 distinct gene products. Target genes of each cell line were stratified into the high-responsiveness group (*Ln*(*IC*50) ≤ 2.36) and the intermediate-responsiveness group (*Ln*(*IC*50) *>* 2.36)^64^.

The Shusi core gene filtering pipeline comprises two sequential steps. First, edges appearing in fewer than 5% of cells within a given cell line were removed to ensure interaction reproducibility. Next, single-cell networks were aggregated, and genes with degree centrality ≥ 250 were retained as high-confidence candidates. For comparative analysis, an equal number of genes were randomly sampled using Python’s random module to generate a background gene list.

### AML patient samples data analysis

Shusi predicted single cell protein networks of all cells cross 27 primary AML specimens. And the single cell network degree (*D_g_*) of each gene was computed by summing all connected edges:

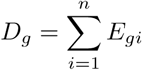

where *E_gi_* represents edges between gene *g* and its interacting partners. To construct high-confidence cell-type-specific networks, we first analyzed monocytic subsets by calculating interaction recurrence rates across all monocytic cells and filtering out interactions appearing in fewer than 5% of cells. We applied identical recurrence-based filtering (5% threshold) to primitive subsets. For identification of monocyte-specific therapeutic targets, we subsequently removed all edges shared with primitive subset networks, thereby isolating uniquely monocytic interactions while maintaining biological relevance through stringent specificity criteria.

### Primary human AML samples

Primary human AML samples were obtained from leukapheresis products of AML patients who had provided written informed consent, with the approval of the local human subject research ethics board at The First Affiliated Hospital of Zhejiang University. The use of the primary human AML samples was reviewed and approved by the Clinical Research Ethics Committee of the First Affiliated Hospital, Zhejiang University School of Medicine.

### Cell culture

Primary human AML cells were stored in a freezing medium composed of 50% Fetal Bovine Serum (FBS; Gibco), 40% Iscove’s Modified Dulbecco’s Medium (IMDM; Gibco) and 10% dimethyl sulfoxide (DMSO; Macklin). The cells were subsequently cryopreserved in liquid nitrogen. Freshly thawed cells were cultured in complete medium at 37^◦^C within a 5% CO_2_ incubator. The complete medium is composed of IMDM (Gibco), 10% FBS (Gibco) and 1% Penicillin/Streptomycin (Yeasen), with the addition of FLT3L, IL-3 and SCF cytokines (Novoprotein), each at a concentration of 10 ng/mL.

### Drug treatment

Freshly thawed primary human AML cells were seeded onto U-bottom 96-well plates at a density of 50 K cells in 150 *µ*L of the complete medium. The cells were then incubated with drugs for 48 hours. All drugs utilized in this study were purchased from MedChemExpress, with each drug being supplied at a stock concentration of 10 mmol/L. A comprehensive list of the drugs is provided in the Supplementary Table 2.

### Immunophenotyping

Immunophenotyping was performed 48 hours after drug treatment. The cells were pelleted and subsequently stained with a panel of flow antibodies, including anti-human CD45 (BD Biosciences; cat. #560566), CD34 (BD Biosciences; cat. #562577), CD117 (BioLegend; cat. #313218), CD11b (BD Biosciences; cat. #555388), CD14 (BD Biosciences; cat. #555397), CD36 (BD Biosciences; cat. #550956), CD64 (BD Biosciences; cat. #563459), CD4 (BD Biosciences; cat. #563028) and CD85k (BioLegend; cat. #333011) at room temperature for 15 minutes, after which they were washed with ice-cold FACS buffer. Following staining, cells were resuspended in FACS buffer containing LIVE/DEAD (Thermo Fisher Scientific; cat. #L34976) and analyzed on Attune NxT (Thermo Fisher Scientific). FCS files were analyzed on FlowJo 10.8.1 (FlowJo). The tSNE analysis of high-dimensional flow data based on dimensionality reduction and automated clustering of concatenated samples was performed in FlowJo using its native platform. A comprehensive list of the flow antibodies is provided in the Supplementary Table 3.

## Data availability

In this study, we used publicly available datasets as detailed in Methods. The specific datasets used are: human cell line dataset available through the Broad Institute’s single-cell portal (SCP542), human cell line deposited in China National GeneBank DataBase (CNGBdb) Sequence Archive (CNSA)(CNP0004330), AML patient samples (GSE232559), and STRING database (https://stringdb-downloads.org/download/protein.links.full.v12.0.txt.gz).

## Code availability

The source code of Shusi is available at GitHub (https://github.com/jiajunyu1999/Shusi).

## Acknowledgements

We thank Bingqian Jiang for helping design the logo of Shusi. We thank for the technical support by the Core Facilities of Liangzhu Laboratory. This work was supported by the Special Funds of the National Natural Science Foundation of China (82450116) and Key R&D Program of Zhejiang (no. 2024SSYS0022). S.P. was supported by the National Natural Science Foundation of China (82470149, 82270157), the Key Research and Development Program of Zhejiang Province (2022C03005), and the Department of Science and Technology of Zhejiang Province (2023R01012).

## Author information

These authors contributed equally: Tianyun Zhang, Jiajun Yu, Shang Lou.

## Authors and Affiliations

**Department of Obstetrics and Gynecology of Sir Run Run Shaw Hospital & Liangzhu Laboratory, Zhejiang University School of Medicine, Hangzhou, China**

Tianyun Zhang, Zhekai Li, & Ning Shen

**Bone Marrow Transplantation Center of The First Affiliated Hospital & Liangzhu Laboratory, Zhejiang University School of Medicine, Hangzhou, China**

Shang Lou, Yue Liang, & Shanshan Pei

**College of Computer Science, Zhejiang University, Hangzhou, China**

Jiajun Yu, Yaozhen Liang & Haishuai Wang

## Contributions

N.S. conceived and guided the study. T.Z. collected data, performed data analysis, and drafted the original manuscript. J.Y., H.W., T.Z., and N.S. designed the model. J.Y. and Y.L. trained the model and performed data analysis. Z.L. and Y.L. collected data and performed data analysis. S.P. and S.L. designed the experiment. S.L. performed the experiments, processed the data and performed data analysis. N.S., S.P., J.Y., S.L.and H.W. revised the manuscript. N.S., S.P., and H.W. supervised the study.

## Corresponding author

Correspondence to Haishuai Wang, Shanshan Pei and Ning Shen

## Ethics declarations

### Competing interests

No authors declare competing interests.

## Notes

### Competing Interest Statement

The authors have declared no competing interest.

### Summary of Updates

We have revised the manuscript results and figures to better clarify our point.

## References

1. Huttlin, E. L. et al. Dual proteome-scale networks reveal cell-specific remodeling of the human interactome. Cell 184, 3022–3040.e28. issn: 00928674. https://linkinghub.elsevier.com/retrieve/pii/S0092867421004463 (2023) (May 2021).

2. Mo, X., et al. Systematic discovery of mutation-directed neo-protein-protein interactions in cancer. Cell 185. Publisher: Elsevier, 1974–1985.e12. issn: 0092-8674, 1097-4172. https://www.cell.com/cell/abstract/S0092-8674(22)00461-5 (2023) (May 26, 2022).

3. Patel, A. S. & Yanai, I. A developmental constraint model of cancer cell states and tumor heterogeneity. Cell 187. Publisher: Elsevier, 2907–2918. issn: 0092-8674, 1097-4172. https://www.cell.com/cell/abstract/S0092-8674(24)00458-6 (2025) (June 6, 2024).

4. Van Galen, P. et al. Single-Cell RNA-Seq Reveals AML Hierarchies Relevant to Disease Progression and Immunity. Cell 176, 1265–1281.e24. issn: 00928674. https://linkinghub.elsevier.com/retrieve/pii/S0092867419300947 (2025) (Mar. 2019).

5. Zeng, A. G. X. et al. A cellular hierarchy framework for understanding heterogeneity and predicting drug response in acute myeloid leukemia. Nature Medicine 28. Publisher: Nature Publishing Group, 1212–1223. issn: 1546-170X. https://www.nature.com/articles/s41591-022-01819-x (2025) (June 2022).

6. González-Silva, L., Quevedo, L. & Varela, I. Tumor Functional Heterogeneity Unraveled by scRNA-seq Technologies. Trends in Cancer 6. Publisher: Elsevier, 13–19. issn: 2405-8033. https://www.cell.com/trends/cancer/abstract/S2405-8033(19)30258-4 (2025) (Jan. 1, 2020).

7. Savage, S. R., et al. Pan-cancer proteogenomics expands the landscape of therapeutic targets. Cell 187. Publisher: Elsevier, 4389–4407.e15. issn: 0092-8674, 1097-4172. https://www.cell.com/cell/abstract/S0092-8674(24)00583-X (2025) (Aug. 8, 2024).

8. Cohen, Y. C. et al. Identification of resistance pathways and therapeutic targets in relapsed multiple myeloma patients through single-cell sequencing. Nature Medicine 27. Number: 3 Publisher: Nature Publishing Group, 491–503. issn: 1546-170X. https://www.nature.com/articles/s41591-021-01232-w (2024) (Mar. 2021).

9. Salcher, S. et al. High-resolution single-cell atlas reveals diversity and plasticity of tissue-resident neutrophils in non-small cell lung cancer. Cancer Cell 40. Publisher: Elsevier, 1503–1520.e8. issn: 1535-6108, 1878-3686. https://www.cell.com/cancer-cell/abstract/S1535-6108(22)00499-8 (2024) (Dec. 12, 2022).

10. Hahn, W. C., et al. An expanded universe of cancer targets. Cell 184. Publisher: Elsevier, 1142–1155. issn: 0092-8674, 1097-4172. https://www.cell.com/cell/abstract/S0092-8674(21)00170-7 (2025) (Mar. 4, 2021).

11. Rood, J. E., Maartens, A., Hupalowska, A., Teichmann, S. A. & Regev, A. Impact of the Human Cell Atlas on medicine. Nature Medicine 28. Number: 12 Publisher: Nature Publishing Group, 2486–2496. issn: 1546-170X. https://www.nature.com/articles/s41591-022-02104-7(2023) (Dec. 2022).

12. Van de Sande, B. et al. Applications of single-cell RNA sequencing in drug discovery and development. Nature Reviews Drug Discovery 22. Number: 6 Publisher: Nature Publishing Group, 496–520. issn: 1474-1784. https://www.nature.com/articles/s41573-023-00688-4 (2025) (June 2023).

13. Vitale, I., Shema, E., Loi, S. & Galluzzi, L. Intratumoral heterogeneity in cancer progression and response to immunotherapy. Nature Medicine 27. Number: 2 Publisher: Nature Publishing Group, 212–224. issn: 1546-170X. https://www.nature.com/articles/s41591-021-01233-9(2025) (Feb. 2021).

14. Van der Weyden, L. et al. Genome-wide in vivo screen identifies novel host regulators of metastatic colonization. Nature 541. Number: 7636 Publisher: Nature Publishing Group, 233–236. issn: 1476-4687. https://www.nature.com/articles/nature20792 (2023) (Jan. 2017).

15. Aibar, S., et al. SCENIC: single-cell regulatory network inference and clustering. Nature Methods 14. Publisher: Nature Publishing Group, 1083–1086. issn: 1548-7105. https://www.nature.com/articles/nmeth.4463 (2025) (Nov. 2017).

16. Bravo González-Blas, C., et al. SCENIC+: single-cell multiomic inference of enhancers and gene regulatory networks. Nature Methods 20. Publisher: Nature Publishing Group, 1355–1367. issn: 1548-7105. https://www.nature.com/articles/s41592-023-01938-4 (2025) (Sept. 2023).

17. Morabito, S., Reese, F., Rahimzadeh, N., Miyoshi, E. & Swarup, V. hdWGCNA identifies co-expression networks in high-dimensional transcriptomics data. Cell Reports Methods 3. Publisher: Elsevier. issn: 2667-2375. https://www.cell.com/cell-reports-methods/abstract/S2667-2375(23)00127-3(2025) (June 26, 2023).

18. Johnson, E. C. B. et al. Large-scale proteomic analysis of Alzheimer’s disease brain and cerebrospinal fluid reveals early changes in energy metabolism associated with microglia and astrocyte activation. Nature Medicine 26. Publisher: Nature Publishing Group, 769–780. issn: 1546-170X. https://www.nature.com/articles/s41591-020-0815-6 (2025) (May 2020).

19. Swarup, V. et al. Identification of Conserved Proteomic Networks in Neurodegenerative Dementia. Cell Reports 31. Publisher: Elsevier. issn: 2211-1247. https://www.cell.com/cell-reports/abstract/S2211-1247(20)30788-9(2025) (June 23, 2020).

20. Cheng, J. et al. ETV7 limits the antiviral and antitumor efficacy of CD8+ T cells by diverting their fate toward exhaustion. Nature Cancer 6. Publisher: Nature Publishing Group, 338–356. issn: 2662-1347. https://www.nature.com/articles/s43018-024-00892-0 (2025) (Feb. 2025).

21. Schäfer, S., et al. scDrugPrio: a framework for the analysis of single-cell transcriptomics to address multiple problems in precision medicine in immune-mediated inflammatory diseases. Genome Medicine 16, 42. issn: 1756-994X. 10.1186/s13073-024-01314-7 (2025) (Mar. 20, 2024).

22. Buccitelli, C. & Selbach, M. mRNAs, proteins and the emerging principles of gene expression control. Nature Reviews Genetics 21. Number: 10 Publisher: Nature Publishing Group, 630–644. issn: 1471-0064. https://www.nature.com/articles/s41576-020-0258-4(2024) (Oct. 2020).

23. Jumper, J., et al. Highly accurate protein structure prediction with AlphaFold. Nature 596. Publisher: Nature Publishing Group, 583–589. issn: 1476-4687. https://www.nature.com/articles/s41586-021-03819-2 (2025) (Aug. 2021).

24. Barabási, A.-L., Gulbahce, N. & Loscalzo, J. Network medicine: a network-based approach to human disease. Nature Reviews Genetics 12. Publisher: Nature Publishing Group, 56–68. issn: 1471-0064. https://www.nature.com/articles/nrg2918 (2025) (Jan. 2011).

25. Zheng, F. et al. Interpretation of cancer mutations using a multiscale map of protein systems. Science 374, eabf3067. issn: 0036-8075, 1095-9203. https://www.science.org/doi/10.1126/science.abf3067 (2025) (Oct. 2021).

26. Li, M. M., et al. Contextual AI models for single-cell protein biology. Nature Methods 21. Publisher: Nature Publishing Group, 1546–1557. issn: 1548-7105. https://www.nature.com/articles/s41592-024-02341-3 (2025) (Aug. 2024).

27. Schaffer, L. V., et al. Multimodal cell maps as a foundation for structural and functional genomics. Nature. Publisher: Nature Publishing Group, 1–10. issn: 1476-4687. https://www.nature.com/articles/s41586-025-08878-3 (2025) (Apr. 9, 2025).

28. Sheinin, R., Sharan, R. & Madi, A. scNET: learning context-specific gene and cell embeddings by integrating single-cell gene expression data with protein–protein interactions. Nature Methods. Publisher: Nature Publishing Group, 1–9. issn: 1548-7105. https://www.nature.com/articles/s41592-025-02627-0 (2025) (Mar. 17, 2025).

29. Xu, K., Hu, W., Leskovec, J. & Jegelka, S. How Powerful are Graph Neural Networks? Feb. 22, 2019. arXiv: 1810.00826[cs]. http://arxiv.org/abs/1810.00826 (2025).

30. Thirunavukarasu, A. J., et al. Large language models in medicine. Nature Medicine 29. Publisher: Nature Publishing Group, 1930–1940. issn: 1546-170X. https://www.nature.com/articles/s41591-023-02448-8 (2025) (Aug. 2023).

31. Kinker, G. S. et al. Pan-cancer single-cell RNA-seq identifies recurring programs of cellular heterogeneity. Nature Genetics 52. Publisher: Nature Publishing Group, 1208–1218. issn: 1546-1718. https://www.nature.com/articles/s41588-020-00726-6 (2024) (Nov. 2020).

32. Zhu, Q. et al. Single cell multi-omics reveal intra-cell-line heterogeneity across human cancer cell lines. Nature Communications 14. Publisher: Nature Publishing Group, 8170. issn: 2041-1723. https://www.nature.com/articles/s41467-023-43991-9 (2024) (Dec. 9, 2023).

33. Szklarczyk, D. et al. The STRING database in 2023: protein–protein association networks and functional enrichment analyses for any sequenced genome of interest. Nucleic Acids Research 51, D638–D646. issn: 0305-1048, 1362-4962. https://academic.oup.com/nar/article/51/D1/D638/6825349 (2025) (D1 Jan. 6, 2023).

34. Wang, X.-W. et al. Assessment of community efforts to advance network-based prediction of protein–protein interactions. Nature Communications 14, 1582. issn: 2041-1723. https://www.nature.com/articles/s41467-023-37079-7 (2024) (Mar. 22, 2023).

35. Reimers, N. & Gurevych, I. Sentence-BERT: Sentence Embeddings using Siamese BERT-Networks Aug. 27, 2019. arXiv: 1908.10084[cs]. http://arxiv.org/abs/1908.10084 (2025).

36. Taylor, R., et al. Galactica: A Large Language Model for Science Nov. 16, 2022. arXiv: 2211.09085[cs]. http://arxiv.org/abs/2211.09085 (2025).

37. Kipf, T. N. & Welling, M. Variational Graph Auto-Encoders Nov. 21, 2016. arXiv: 1611.07308[stat]. http://arxiv.org/abs/1611.07308 (2025).

38. Gingras, A.-C., Gstaiger, M., Raught, B. & Aebersold, R. Analysis of protein complexes using mass spectrometry. Nature Reviews Molecular Cell Biology 8. Publisher: Nature Publishing Group, 645–654. issn: 1471-0080. https://www.nature.com/articles/nrm2208 (2025) (Aug. 2007).

39. Kim, M. et al. A protein interaction landscape of breast cancer. Science 374, eabf3066. issn: 0036-8075, 1095-9203. https://www.science.org/doi/10.1126/science.abf3066 (2025) (Oct. 2021).

40. Ghandi, M., et al. Next-generation characterization of the Cancer Cell Line Encyclopedia. Nature 569. Publisher: Nature Publishing Group, 503–508. issn: 1476-4687. https://www.nature.com/articles/s41586-019-1186-3 (2025) (May 2019).

41. Yang, W. et al. Genomics of Drug Sensitivity in Cancer (GDSC): a resource for therapeutic biomarker discovery in cancer cells. Nucleic Acids Research 41, D955–D961. issn: 0305-1048, 1362-4962. http://academic.oup.com/nar/article/41/D1/D955/1059448/Genomics-of-Drug-Sensitivity-in-Cancer-GDSC-a (2025) (D1 Nov. 22, 2012).

42. Beneyto-Calabuig, S. et al. Clonally resolved single-cell multi-omics identifies routes of cellular differentiation in acute myeloid leukemia. Cell Stem Cell 30, 706–721.e8. issn: 19345909. https://linkinghub.elsevier.com/retrieve/pii/S1934590923001194 (2025) (May 2023).

43. Lasry, A. et al. An inflammatory state remodels the immune microenvironment and improves risk stratification in acute myeloid leukemia. Nature Cancer 4. Publisher: Nature Publishing Group, 27–42. issn: 2662-1347. https ://www.nature.com/articles/s43018-022-00480-0 (2025) (Jan. 2023).

44. Tyner, J. W., et al. Functional genomic landscape of acute myeloid leukaemia. Nature 562. Publisher: Nature Publishing Group, 526–531. issn: 1476-4687. https://www.nature.com/articles/s41586-018-0623-z (2025) (Oct. 2018).

45. Pei, S. et al. A novel type of monocytic leukemia stem cell revealed by the clinical use of venetoclax-based therapy. Cancer discovery 13, 2032–2049. issn: 2159-8274. https://www.ncbi.nlm.nih.gov/pmc/articles/PMC10527971/ (2025) (Sept. 6, 2023).

46. Pei, S. et al. Monocytic Subclones Confer Resistance to Venetoclax-Based Therapy in Patients with Acute Myeloid Leukemia. Cancer Discovery 10, 536–551. issn: 2159-8274, 2159-8290. https://aacrjournals.org/cancerdiscovery/article/10/4/536/2403/Monocytic-Subclones-Confer-Resistance-to (2025) (Apr. 1, 2020).

47. Ellegast, J. M. et al. Unleashing cell-intrinsic inflammation as a strategy to kill AML blasts. Cancer discovery 12, 1760–1781. issn: 2159-8274. https://www.ncbi.nlm.nih.gov/pmc/articles/PMC9308469/ (2025) (July 6, 2022).

48. Wang, B. et al. Comprehensive characterization of IFN signaling in acute myeloid leukemia reveals prognostic and therapeutic strategies. Nature Communications 15. Publisher: Nature Publishing Group, 1821. issn: 2041-1723. https://www.nature.com/articles/s41467-024-45916-6 (2025) (Feb. 28, 2024).

49. Nwosu, G. O., Ross, D. M., Powell, J. A. & Pitson, S. M. Venetoclax therapy and emerging resistance mechanisms in acute myeloid leukaemia. Cell Death & Disease 15. Publisher: Nature Publishing Group, 1–11. issn: 2041-4889. https://www.nature.com/articles/s41419-024-06810-7 (2025) (June 12, 2024).

50. Luck, K., et al. A reference map of the human binary protein interactome. Nature 580. Publisher: Nature Publishing Group, 402–408. issn: 1476-4687. https://www.nature.com/articles/s41586-020-2188-x (2025) (Apr. 2020).

51. Kipf, T. N. & Welling, M. Semi-Supervised Classification with Graph Convolutional Networks Feb. 22, 2017. arXiv: 1609.02907[cs]. http://arxiv.org/abs/1609.02907 (2025).

52. Hamilton, W. L., Ying, R. & Leskovec, J. Inductive Representation Learning on Large Graphs Sept. 10, 2018. arXiv: 1706.02216[cs]. http://arxiv.org/abs/1706.02216 (2025).

53. Veličković, P., et al. Graph Attention Networks Feb. 4, 2018. arXiv: 1710.10903[stat]. http://arxiv.org/abs/1710.10903 (2025).

54. Brody, S., Alon, U. & Yahav, E. How Attentive are Graph Attention Networks? Jan. 31, 2022. arXiv: 2105.14491[cs]. http://arxiv.org/abs/2105.14491 (2025).

55. Paszke, A. Pytorch: An imperative style, high-performance deep learning library. *arXiv preprint arXiv:1912.01703* (2019).

56. Fey, M. & Lenssen, J. E. Fast graph representation learning with PyTorch Geometric. *arXiv preprint arXiv:1903.02428* (2019).

57. Kingma, D. Adam: a method for stochastic optimization in Int Conf Learn Represent (2014).

58. Oughtred, R. et al. The ¡span style=”font-variant:small-caps;”¿BioGRID¡/span¿ database: A comprehensive biomedical resource of curated protein, genetic, and chemical interactions. Protein Science 30, 187–200. issn: 0961-8368, 1469-896X. https://onlinelibrary.wiley.com/doi/10.1002/pro.3978 (2025) (Jan. 2021).

59. Schweppe, D. K., Huttlin, E. L., Harper, J. W. & Gygi, S. P. BioPlex Display: An Interactive Suite for Large-Scale AP–MS Protein–Protein Interaction Data. Journal of Proteome Research 17. Publisher: American Chemical Society, 722–726. issn: 1535-3893. 10.1021/acs.jproteome.7b00572 (2025) (Jan. 5, 2018).

60. Drew, K., Wallingford, J. B. & Marcotte, E. M. hu.MAP 2.0: integration of over 15,000 proteomic experiments builds a global compendium of human multiprotein assemblies. Molecular Systems Biology. https://www.embopress.org/doi/10.15252/msb.202010016 (2025) (May 11, 2021).

61. Hein, M. Y. et al. A Human Interactome in Three Quantitative Dimensions Organized by Stoichiometries and Abundances. Cell 163, 712–723. issn: 00928674. https://linkinghub.elsevier.com/retrieve/pii/S0092867415012702 (2025) (Oct. 2015).

62. Hao, Y. et al. Dictionary learning for integrative, multimodal and scalable single-cell analysis. Nature Biotechnology. 10.1038/s41587-023-01767-y (2023).

63. Gaulton, A., et al. ChEMBL: a large-scale bioactivity database for drug discovery. Nucleic Acids Research 40, D1100–D1107. issn: 0305-1048. https://www.ncbi.nlm.nih.gov/pmc/articles/PMC3245175/ (2025) (Database issue Jan. 2012).

64. Joo, M. et al. A Deep Learning Model for Cell Growth Inhibition IC50 Prediction and Its Application for Gastric Cancer Patients. International Journal of Molecular Sciences 20, 6276. issn: 1422-0067. https://www.mdpi.com/1422-0067/20/24/6276 (2025) (Dec. 12, 2019).

